# The impact of sparsity in low-rank recurrent neural networks

**DOI:** 10.1101/2022.03.31.486515

**Authors:** Elizabeth Herbert, Srdjan Ostojic

## Abstract

Neural population dynamics are often highly coordinated, allowing task-related computations to be understood as neural trajectories through low-dimensional subspaces. How the network connectivity and input structure give rise to such activity can be investigated with the aid of low-rank recurrent neural networks, a recently-developed class of computational models which offer a rich theoretical framework linking the underlying connectivity structure to emergent low-dimensional dynamics. This framework has so far relied on the assumption of all-to-all connectivity, yet cortical networks are known to be highly sparse. Here we investigate the dynamics of low-rank recurrent networks in which the connections are randomly sparsified, which makes the network connectivity formally full-rank. We first analyse the impact of sparsity on the eigenvalue spectrum of low-rank connectivity matrices, and use this to examine the implications for the dynamics. We find that in the presence of sparsity, the eigenspectra in the complex plane consist of a continuous bulk and isolated outliers, a form analogous to the eigenspectra of connectivity matrices composed of a low-rank and a full-rank random component. This analogy allows us to characterise distinct dynamical regimes of the sparsified low-rank network as a function of key network parameters. Altogether, we find that the low-dimensional dynamics induced by low-rank connectivity structure are preserved even at high levels of sparsity, and can therefore support rich and robust computations even in networks sparsified to a biologically-realistic extent.

**Author summary:** In large networks of neurons, the activity displayed by the population depends on the strength of the connections between each neuron. In cortical regions engaged in cognitive tasks, this population activity is often seen to be highly coordinated and low-dimensional. A recent line of theoretical work explores how such coordinated activity can arise in a network of neurons in which the matrix defining the connections is constrained to be mathematically low-rank. Until now, this connectivity structure has only been explored in fully-connected networks, in which every neuron is connected to every other. However, in the brain, network connections are often highly sparse, in the sense that most neurons do not share direct connections. Here, we test the robustness of the theoretical framework of low-rank networks to the reality of sparsity present in biological networks. By mathematically analysing the impact of removing connections, we find that the low-dimensional dynamics previously found in dense low-rank networks can in fact persist even at very high levels of sparsity. This has promising implications for the proposal that complex cortical computations which appear to rely on low-dimensional dynamics may be underpinned by a network which has a fundamentally low-rank structure, albeit with only a small fraction of possible connections present.

## Introduction

Neural recordings in animals performing cognitive tasks have revealed that individual neurons ubiquitously display a high degree of coordination. When viewed in the activity state space, in which each each axis represents the firing rate of one unit, the trajectories of neural activity are typically confined to low-dimensional subspaces [1–6]. The resulting latent dynamics have been proposed to underpin complex cortical computations [7]. However, how the inputs to the network and the connectivity between the individual units shape such low-dimensional activity remains a prominent question.

A recently developed class of models, recurrent networks with low-rank connectivity, provide a tractable theoretical framework for addressing this question and unravelling the relationship between connectivity structure, low-dimensional dynamics and the resulting computations [8–17]. One important limitation is, however, that these models often assume a dense connectivity structure in which every neuron shares synapses with every other. In contrast, cortical networks exhibit a high degree of sparsity in their connectivity, meaning that each neuron receives inputs from only a fraction of its neighbors [18–21]. Since sparse matrices are typically full-rank, an important question is whether and how the results obtained in the study of low-rank recurrent networks apply to sparse connectivity structures.

Here we investigate how the dynamics of low-rank recurrent networks are impacted by increasing degrees of sparsity. We start by analysing the eigenvalue spectra of low-rank connectivity matrices in which sparsity is imposed by removing a random fraction of entries. Such matrices are full rank, but we find that the corresponding eigenspectra consist of a continuous bulk and isolated outliers, and are therefore analogous to low-rank matrices superposed with a random, full rank component [8, 22, 23]. We show that both the radius of the eigenvalue bulk and the outliers can be estimated analytically. We then use these results to compare the dynamics of sparsified low-rank networks to those of densely connected low-rank networks with a full rank random component. Altogether we found that the low-dimensional dynamics generated by a low-rank connectivity structure are highly resistant with respect to sparsity and therefore provide a robust substrate for implementing computations in networks with biologically realistic connectivity.

## Results

### Network connectivity

We study recurrent networks of *N* firing rate units, following the classical formalism in [24]. The dynamics of each individual unit *i* evolve as:

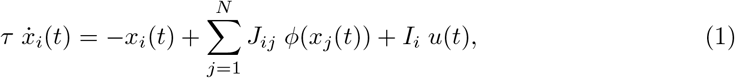

where *x_i_* describes the total input current to each unit, *τ* is the time constant of the dynamics, is the synaptic weights from unit *j* to unit *i* and *ϕ*(·) is a non-linear transfer function that we take to be the hyperbolic tangent. Each unit can also receive a time-dependent input current of magnitude *u*(*t*) via a feedforward weight vector **I** ={*I_i_*}_*i*=1…*N*_. The set of synaptic weights *J_ij_* are stored in a connectivity matrix **J**, for which we consider two forms. We begin by considering full-rank Gaussian connectivity, introducing sparsity into the synaptic weights and establishing the ways in which sparse Gaussian networks differ from their fully-connected counterparts. We then constrain the connectivity to be low-rank and sparsify as before, examining how the impact of sparsity in such networks both parallels and contrasts with that of the Gaussian case.

The full-rank Gaussian networks are defined by a matrix **J** with entries independently distributed as

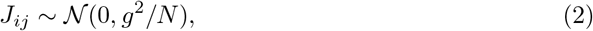

where *g* controls the variance of the matrix entries and thus the strength of the coupling.

For the low-rank networks, we consider the simplest case of a rank-one matrix **P**, constructed as in previous work [8] as a rescaled outer product of two *N*-dimensional random connectivity vectors **m** = {*m_i_*}_*i*=1…*N*_ and **n** = {*n_i_*}_*i*=1…*N*_ such that

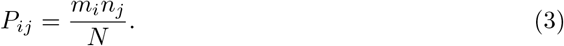

This ensures that all columns of **P** are linearly dependent and proportional to **m**. The individual entries *m_i_* and *n_i_* of the connectivity vectors are drawn independently for each *i* from a joint Gaussian distribution with mean 0 and covariance matrix Σ:

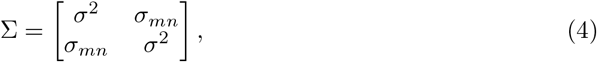

where *σ*^2^ is the variance of both connectivity vectors which controls the overall strength of the coupling, and *σ_mn_* is the covariance between them (see Methods). In the large *N* limit, this covariance becomes equivalent to the degree of overlap between **m** and **n**, given by the normalised scalar product:

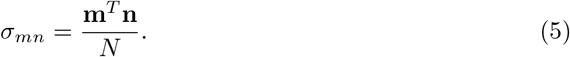

The covariance *σ_mn_* plays a critical role in the stability of the dynamics of the rank-one network due to its influence on the matrix eigenvalues, as will be seen in the following sections. The variance *σ*^2^ likewise gains a critical influence as soon as the matrix becomes sparse. These two key parameters controlling the connectivity, the variance and covariance of the connectivity vectors, will therefore become paramount in the later analysis of the dynamics.

### Eigenvalues of connectivity matrices

The dynamics of recurrent networks are strongly influenced by the eigenspectrum of their connectivity matrix. Regardless of network structure, the dynamics always possess a trivial fixed point at zero, since we take the transfer function *ϕ*(·) to be the hyperbolic tangent, and tanh(0) = 0. The stability of this zero fixed point is determined by the magnitude of the eigenvalue with largest real part, λ*^max^*. Since *ϕ*′(0) = 1, the stability matrix at zero reduces to *S_ij_* = *J_ij_* – *δ_ij_*, so the fixed point at zero becomes unstable when the largest eigenvalue of the connectivity matrix **J** surpasses unity. As soon as this occurs, non-trivial dynamics can emerge. To understand the impact of sparsity on network dynamics we will therefore analyse the changes in the network eigenspectra, and will place particular focus on λ*^max^*, the eigenvalue with maximum real part.

### Eigenvalues of sparsified full-rank networks

We first consider the impact of sparsity on the eigenspectra of the full-rank, random connectivity matrices (Eq. 2). Work in random matrix theory has demonstrated that the eigenspectra of such matrices are described by Girko’s circular law [25]: for a matrix with entries independently distributed with mean zero and variance *Var*, the eigenvalues converge in the limit of *N* → ∞ to a uniform distribution within a disk of radius 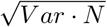 centred at the origin. This result is universal in the sense that it holds for any distribution with finite variance. For the Gaussian networks **J** introduced in (Eq. 2) with *Var* = *g*^2^/*N*, the eigenvalues are therefore uniformly distributed on a circular disk of radius approximated by *g*. Since the distribution is circular, the radius of this disk is equivalent in the large *N* limit to λ*^max^*, the eigenvalue with maximum real part, so we refer to both by the spectral radius *R*.

To sparsify the matrix, we simply choose a fraction *s* ∈ [0, 1] of connections to set to zero at random. This is achieved by multiplying the original matrix **J** elementwise with a binary matrix **X**, where *X_ij_* are drawn independently from a Bernouilli distribution 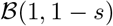, forming a sparse matrix 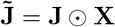 characterised by a degree of sparsity *s* (Fig 1A). Due to the presence of zeros, the entries of 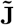 have a lower variance than those of **J**, but remain independently distributed. Because of this property of independence, we expect the universality result of the circular law for iid matrices to hold [25]. Indeed, we find that the eigenvalues of the sparse matrix 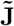 continue to distribute uniformly on a disk for which the spectral radius is described by 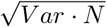, where *Var* is now the variance of the sparse matrix elements 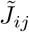. This variance can be calculated (see Methods) as (1 – *s*) *g*^2^/*N*, giving a spectral radius of:

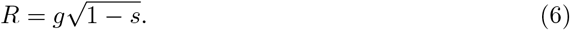

**Fig 1.**
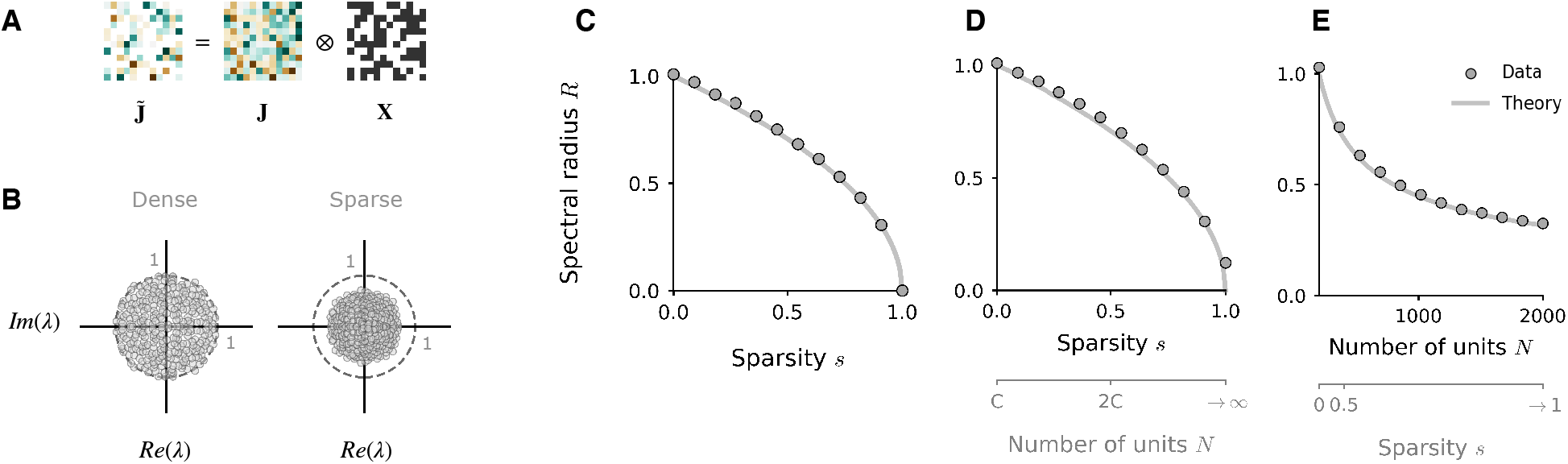
Influence of sparsity on the eigenspectra of full-rank networks. **A**: Illustration of how sparsity is imposed in the connectivity matrix, where the degree of sparsity is *s* = 0.5. **B**: Complex eigenspectra of full rank, Gaussian connectivity matrices of finite size (*N* = 300, *g* = 1) in the dense case (left) and with a sparsity of 0.5 (right). The dashed line plots the unit circle. **C, D, E**: Reduction of spectral radius *R* as a function of sparsity in a full-rank matrix **J** constructed as in (Eq. 2) with connection strength *g* = 1. In **C**, sparsity is imposed as a fraction of total connections removed (N = 1000). In **D** and **E** sparsity is imposed by fixing the number of outgoing connections to *C* = 200 and increasing *N*. Dots: mean empirical spectral radius, measured as the largest absolute value of all eigenvalues, over 50 instances. Solid lines: theoretical prediction.

Increasing the degree of sparsity *s* in a full-rank Gaussian network thus monotonically reduces the radius of the disk on which the eigenvalues distribute. In Fig 1C we demonstrate the correspondence of the prediction in (Eq. 6) to the empirical spectral radius, measured as the largest eigenvalue in the spectrum of a finite-sized Gaussian matrix sparsified in the manner described above.

An alternative means of sparsifying the matrix is to set to zero a fixed number of outgoing connections *C* per unit, while increasing the total number of units *N*, where *C* ≤ *N*. This is often a regime of interest in neuroscientific work [26]. The network is now defined by a degree of sparsity *s* = 1 – *C*/*N*, where *C*/*N* is the fraction of non-zero connections per unit. The impact of sparsifying the matrix in this manner is equivalent to the previous case, where the theoretical spectral radius is now given by 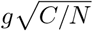 (Fig 1D) and taking the degree of sparsity to one now corresponds to taking the number of units *N* to infinity (Fig 1E).

### Eigenvalues of sparsified rank-one networks

We now turn to the impact of sparsity on the eigenspectra of rank-one matrices. Fully-connected rank-one matrices have only one potentially-nonzero eigenvalue, formed from the scalar product of the corresponding left and right eigenvectors. For the matrix **P** defined in (Eq. 3), the right and left eigenvectors are the connectivity vectors **m** and **n**, and the corresponding eigenvalue is located at **m**^*T*^**n**/*N* on the real axis, which is equivalent in the large *N* limit to the overlap *σ_mn_* between the connectivity vectors.

When sparsity is introduced to the rank-one structure, forming a sparse matrix 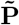, this matrix is now formally full rank and possesses *N* potentially-nonzero eigenvalues. However, we empirically observed that the eigenspectrum splits into two distinct components. The eigenvalue associated with the rank-one structure remains distinct in the spectrum, since such structure persists as a backbone to the connectivity. We refer to this structural eigenvalue as the *outlier*. At the same time, the full-rank perturbation introduced by the sparsity induces additional eigenvalues with nonzero real and imaginary parts which distribute about the origin on a disk with non-uniform density (Fig 2A). We refer to this set of additional non-zero eigenvalues induced by sparsity as the *bulk*. For the sparsified rank-one matrix 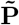, it is now these two components of the spectrum, the bulk and the outlier, that together contribute to the dynamics. Understanding the impact of sparsity on the dynamics therefore reduces to understanding how each component is modified by sparsity. We now address each in turn.

**Fig 2.**
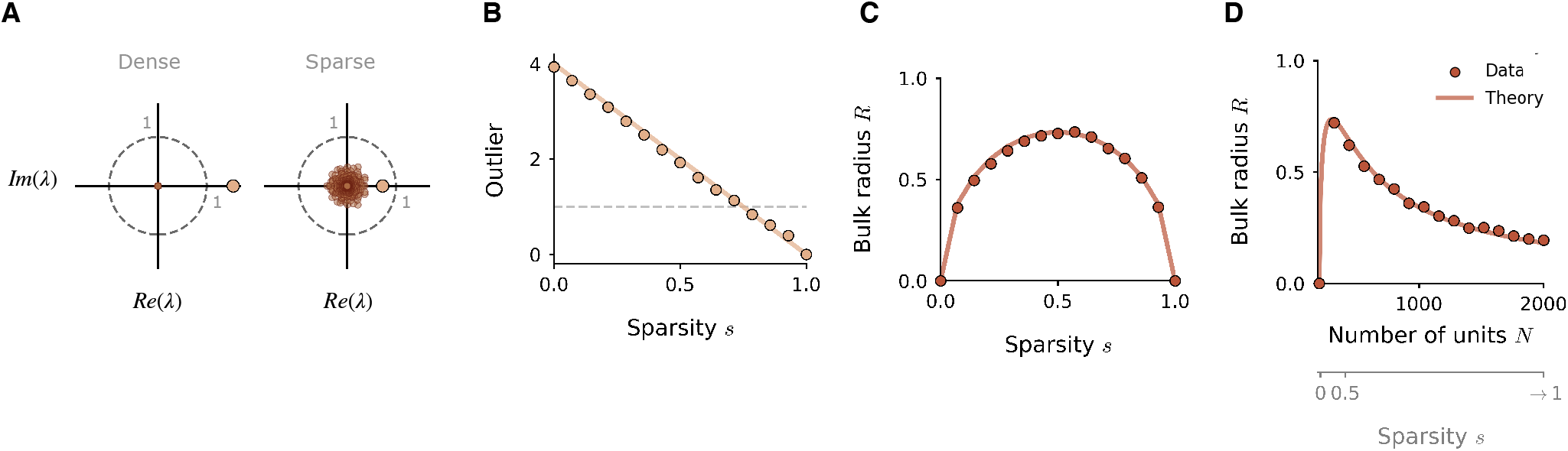
Influence of sparsity on the eigenspectra of rank-one networks. **A**: Eigenspectra of rank-one connectivity matrices of finite size, in the dense case (left) and under a sparsity of 0.5 (right). The matrix **P** is constructed as in (Eq. 3), with parameters *σ*^2^ = 16, *σ_mn_* = 1.44 and *N* = 300. Under sparsity, the outlier (gold) is reduced and the bulk distribution (brown) emerges. The dashed line plots the unit circle. **B, C**: Impact of sparsity on two key features of the eigenspectrum of finite-size rank-one networks: **B**, the outlier λ_1_, and **C**, the spectral radius of the bulk distribution. The outlier is eventually reduced below the instability boundary of λ_1_ = 1, dashed line. Sparsity is imposed as a fraction of total connections removed; *σ*^2^ = 16, *σ_mn_* = 4 and *N* = 1000. **D**: Same as in **C** but for sparsity imposed by fixing *C* = 200 non-zerro connections and increasing *N*; bulk radius is plotted as a function of *N*. Dots: empirical measurements of outlier and bulk radius. Solid lines: theoretical prediction.

Firstly, the outlier λ_1_ is reduced monotonically by sparsity. It can be shown that in the large *N* limit, the right connectivity vector **m** remains a right-eigenvector of 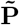, and yet a fraction *s* of matrix entries are now zero; this means that the factor by which the outlier is reduced is 1 – *s* (see Methods). The outlier therefore lies at (1 – *s*) **m**^*T*^**n**/*N* on the real axis, and is drawn in towards the origin as the degree of sparsity approaches 1 (Fig 2B). In the large *N* limit, this value is equivalent to

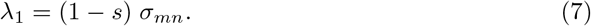

Secondly, we wish to characterise the radius of the bulk distribution. Although the distribution of eigenvalues in the bulk is non-uniform (Fig 3A), it continues to be circular, and we thus hypothesise that the universality result for the radius [25] still holds. This would allow us to characterise the bulk radius directly using the variance of matrix elements, in a similar manner to the Gaussian matrix. To test this, we derive the variance of the elements not of the entire sparse matrix 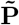, but of a new matrix 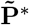 which possesses solely the eigenvalues in the bulk distribution. We remove the eigenvalue outlier from the spectrum as in [22, 23, 27], constructing a new matrix 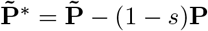 as a linear combination of the dense matrix **P** and the original sparse matrix 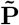. The connectivity vector **m** is also an eigenvector of 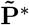, but now with a zero eigenvalue. The distribution of the remaining eigenvalues of 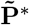 is identical to those in 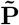, but with the outlier λ_1_ removed. By deriving the variance of the elements of 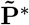 (see Methods), we obtain:

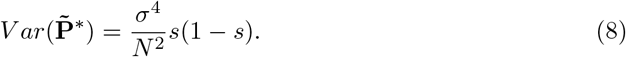

**Fig 3.**
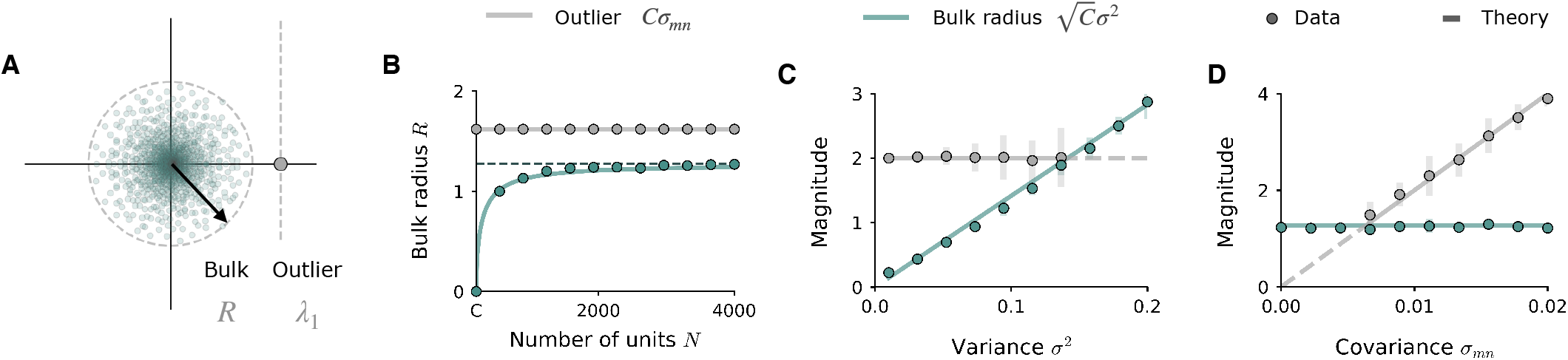
Key features of rank-one eigenspectrum become independent of *N* in the high sparsity limit. **A**: Illustration of the spectral radius *R* of the bulk distribution induced by sparsity, and the outlier λ_1_ inherited from the rank-one structure. Dots: eigenvalues of matrix. Dashed lines: theoretical predictions, with *C* = 200, *N* = 2000, *σ*^2^ = 0.09, and *σ_mn_* = 0.008. **B**: Bulk radius as a function of sparsity imposed by fixing *C* = 200 and increasing *N*, for the rescaled matrix *P_ij_* = *m_i_n_j_* with *σ*^2^ = 0.09 and *σ_mn_* = 0.008. The outlier *Cσ_mn_* is now independent of *N*. The bulk radius converges towards 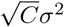 (grey dashed line) as sparsity increases. **D**: Outlier and bulk radius as a function of the variance of the connectivity vectors, while the covariance is fixed (*σ_mn_* = 0.01). **E**: Outlier and bulk radius as a function of the covariance, while the variance is fixed (*σ*^2^ = 0.09). Empirical values are displayed as mean (dots) and standard deviation (bars, 10 repeats) of the eigenvalue with largest absolute magnitude (bulk) and real part (outlier), while the outlier is still distinguished from the bulk. When the outlier is smaller than the bulk, its location cannot be measured empirically. Lines: theoretical predictions at empirically measurable (solid) and unmeasurable (dashed) locations. Parameters: *C* = 200, *N* = 1200, resulting in *s* = 0.8.

The theoretical bulk radius given by the circular law is therefore:

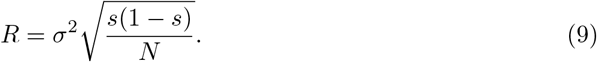

We find that this expression accurately describes the radius of the bulk distribution measured empirically in finite-size networks (Fig 2C). When sparsity is imposed by setting a fixed number of connections *C* and increasing *N*, these quantities can simply be redefined in terms of *C* and *N* by substituting *s* = 1 – *C*/*N*. We thus obtain the outlier as:

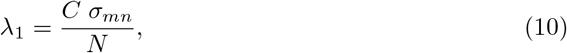

and the radius of the bulk as:

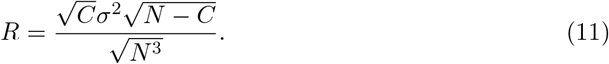

In contrast to the previous case of Gaussian networks, the radius of the bulk distribution which emerges in rank-one networks therefore scales non-monotonically with sparsity, first increasing to its maximum extent at *s* = 0.5, then reducing once more into the origin as the degree of sparsity approaches 1 (Fig 2C, D).

### High sparsity limit

With the rank-one connectivity defined with a 1/*N* scaling as in (Eq. 3), when *C* is fixed and *N* is taken to ∞, both the bulk radius *R* and the outlier λ_1_ are reduced to zero. In order to have a non-vanishing eigenspectrum and ensure that the bulk radius and the outlier remain finite in the limit of *N* → ∞, we turn to a rescaled version of the connectivity matrix *P_ij_*. By removing the weight scaling by *N* and considering simply the matrix

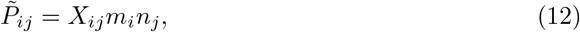

the outlier is now constant with respect to sparsity:

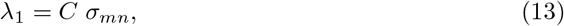

and the radius of the bulk becomes:

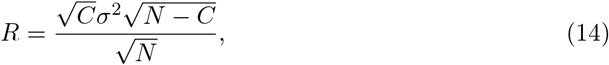

which approaches 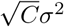 in the limit of high sparsity as *N* is taken to infinity at finite *C* (Fig 3B). Thus in this rescaled network, for a given number of connections per neuron *C* and for a high level of sparsity *C* ≪ *N*, the radius of the bulk distribution depends only on the variance *σ*^2^ of the connectivity vectors (Fig 3C), and the location of the structural eigenvalue outlier depends only on their covariance *σ_mn_* (Fig 3D). This decoupling from *N* allows us to understand the dynamics of networks situated in the high-sparsity regime solely in terms of the two variables characterising the rank-one connectivity, the variance *σ*^2^ and the covariance *σ_mn_* of the connectivity vectors.

### Dynamics of sparsified rank-one networks

Having characterised the eigenspectrum of the sparsified rank-one matrix, we now turn to the insights we can extract about the dynamics. We have shown that the eigenspectrum of the sparse rank-one matrix is comprised of two distinct components, the outlier and the bulk distribution, which are under the independent control of two key parameters defining the network connectivity. Moreover, we have shown that in the large *N* limit, the spectral radius of the bulk distribution can be characterised by Girko’s circular law, in the same manner as the circular disk of eigenvalues characteristic of a full-rank Gaussian matrix (Fig 1). This leads us to hypothesize an equivalence between the dynamics of a sparsified rank-one network and those of a dense rank-one network with an added full-rank, Gaussian component, which also give rise to an outlier and an eigenvalue disk into the spectrum [8]. In what follows, we therefore explore the extent to which the dynamics of sparsified rank-one networks resemble dense rank-one networks with additional random connectivity, and highlight the aspects in which they are unique.

To preface our interpretation of the dynamics, we briefly summarise the behaviour of low-rank recurrent networks in the dense case [8, 12, 13]. In general, a network of rank *R* gives rise to dynamics embedded in an *R*-dimensional subspace spanned by the right connectivity vectors, with an additional dimension introduced by each addition of external input along a given vector **I**. Recent work has demonstrated that the trajectories within this subspace can be reduced to a mean-field description of a small number of interacting latent variables [12, 13]. For the rank-one networks that we consider here, the activation of each unit *x_i_* can be described by:

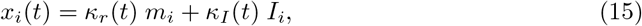

where **I** is the vector along which the network receives an external input. The latent variables *κ_r_*(*t*) and *κ_I_*(*t*) define the projection of the population activity **x** onto the vectors **m** and **I** respectively. The population activity therefore spans the plane formed by the vectors **m** and **I**, and reduces to a one-dimensional trajectory along the vector **m** in the absence of input. Whether or not activity is generated along **m** is determined by the total recurrent input *κ_rec_*, given by (see Methods):

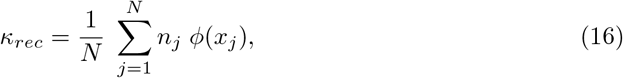

which represents the overlap of the network activity *ϕ*(**x**) with the left connectivity vector **n**. A non-zero value of *κ_rec_* - and thus non-trivial equilibrium dynamics structured along **m** - can only arise if the connectivity vector **n** has a non-zero overlap *σ_nI_* with the input vector **I** (input-driven dynamics) or a non-zero overlap *σ_nm_* with the connectivity vector **m** (autonomous dynamics).

Here, we address the impact of sparsity on the degree of structure present in the input-driven and autonomous network dynamics. We focus on rank-one networks in the high-sparsity limit (Eq. 12) where the number of non-zero connections *C* is fixed independently of *N*, in which case the magnitude of the outlier and bulk distribution become independent of *N* and remain finite at high sparsities. We first fix both the outlier and the bulk below unity, and consider the network response to an external feedforward input (Fig. 4). We then turn to the autonomous dynamics that arise when both the outlier and bulk are above the instability, and study the dynamical landscape formed from the interaction between the two components (Fig. 5).

**Fig 4.**
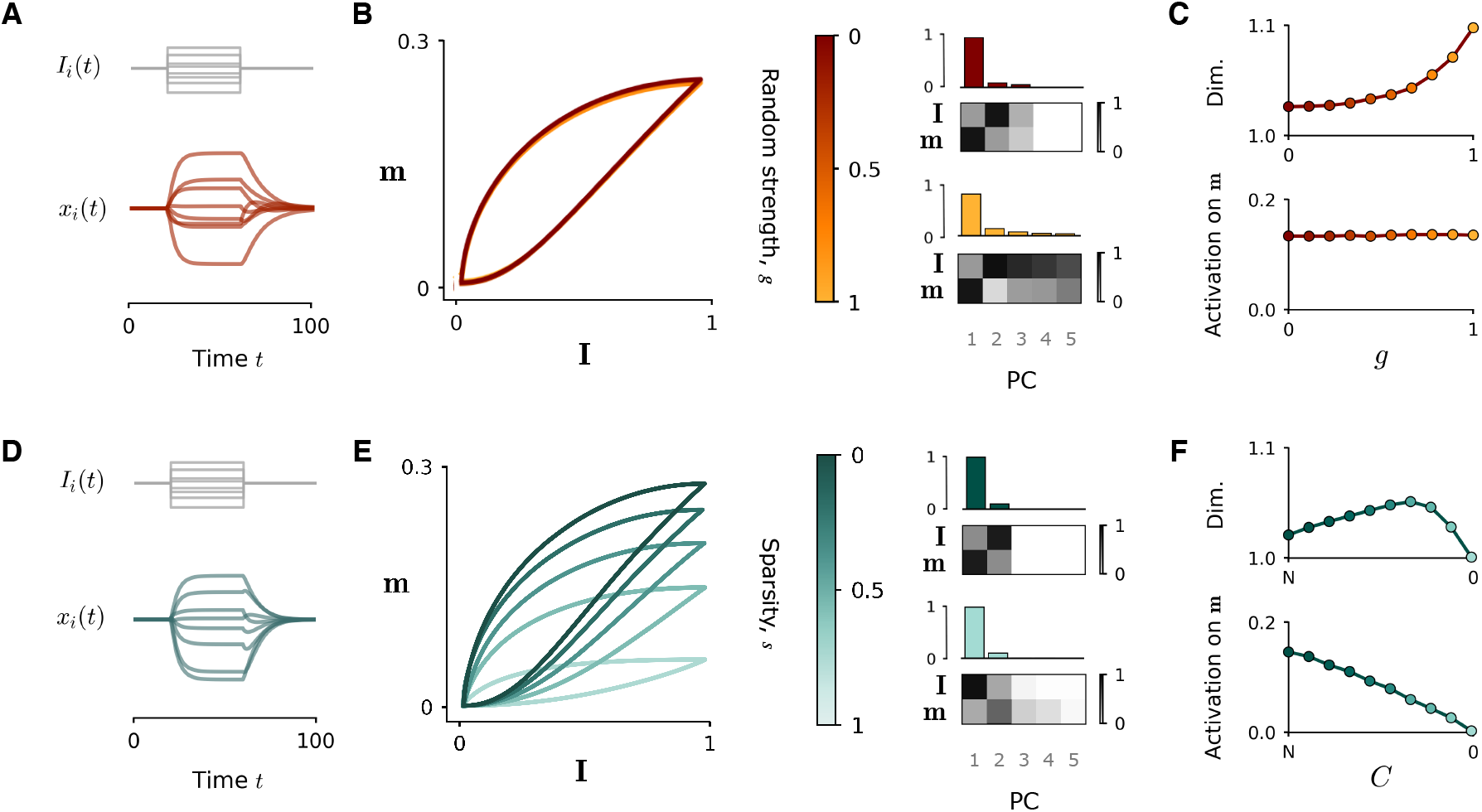
Impact of sparsity on input-driven dynamics. Network responses to a step input current along a random vector **I**. Top row: network consists of a dense rank-one component 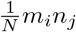 and a full-rank, random component of variance *g*^2^/*N*; the random strength *g* is progressively increased. Bottom row: network consists only of rank-one component *P_ij_* = *m_i_n_j_*; the sparsity is progressively increased by decreasing the number of non-zero connections *C*. Both the bulk and outlier in the eigenspectrum lie below unity, but the input vector **I** partially overlaps with **n** (*σ_nI_* = 0.2). **A, D**: Temporal dynamics of the network during step input (**A**: *g* = 0.8; **D**: *s* = 0.8). Top: samples of input timeseries *I_i_u*(*t*). Bottom: samples of network activations *x_i_*. **B, E**: Left: input-driven population trajectories projected onto the plane defined by the right connectivity vector **m** and input vector **I**, as random strength (resp. sparsity) is progressively increased. Right: principal component analysis (PCA) of each trajectory, showing the fraction of variance explained by the first three components (upper panels) and the correlation between first three principal components and the vectors **I** and **m** (lower panels). **C, F**: Top: Dimensionality of network trajectories quantified by the participation ratio 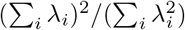, where λ*_i_* are the eigenvalues of the covariance matrix of activations. Bottom: Projection *κ_r_* of network activation **x** onto right connectivity vector **m**. The mean value for both quantities is taken over 50 simulations for each value of *C* and *g*. Parameters for all graphs: *N* = 2000, *σ*^2^ = 0.015, *σ_mn_* = 0.

**Fig 5.**
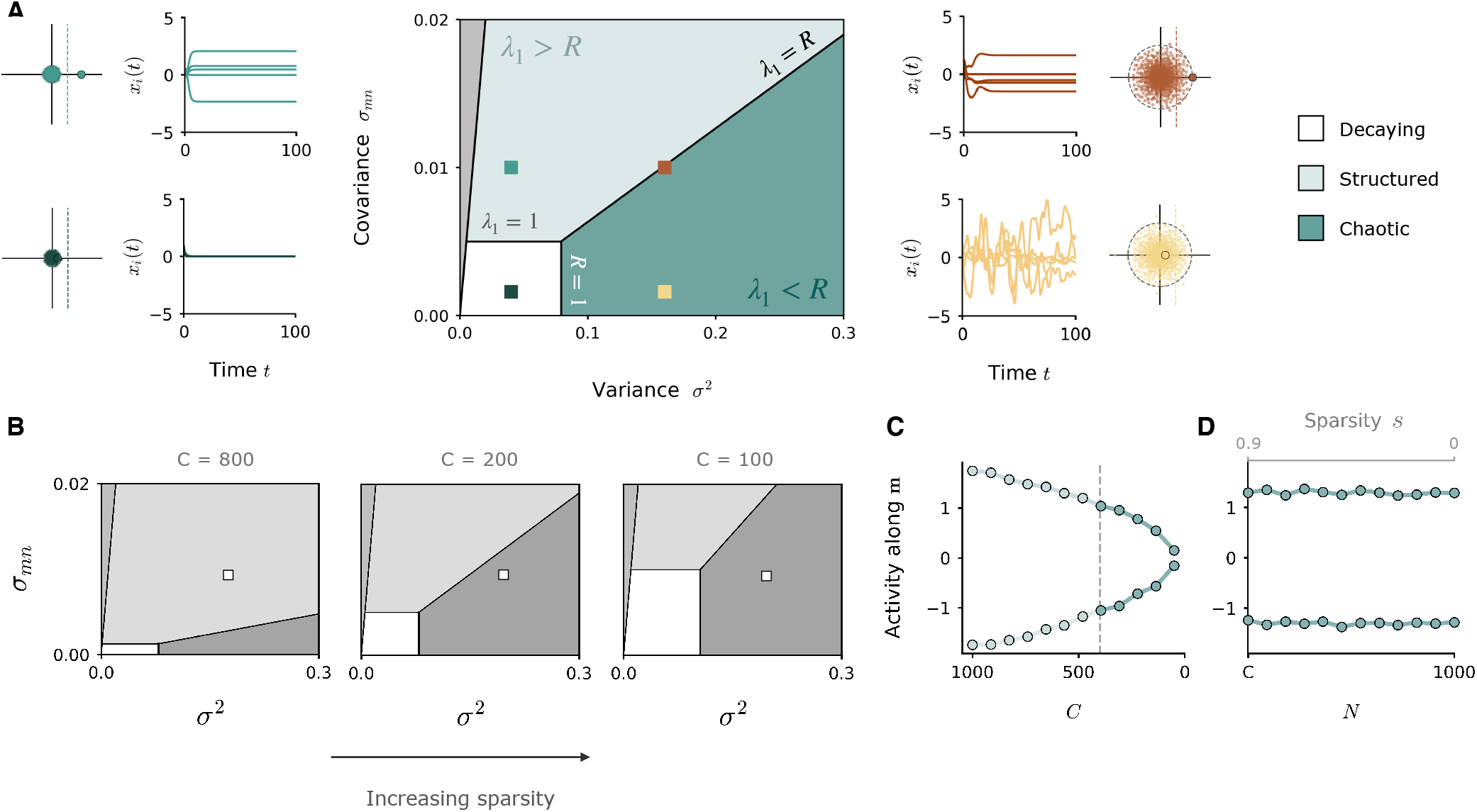
Dynamical regimes of autonomous network activity at high sparsity. **A**: Dynamics of a sparsified rank one network in the high-sparsity regime where *P_ij_* = *m_i_n_j_* and the number of non-zero connections *C* is fixed. The variance *σ*^2^ and covariance *σ_mn_* of the connectivity vectors respectively control the bulk radius and outlier of the eigenvalue distribution. Centre: Phase diagram of dynamical regimes in the variance-covariance plane, for *C* = 200 and *N* = 1000. The transition from structured to chaotic activity occurs when the bulk radius surpasses the location of the outlier. Side panels: samples of autonomous dynamics of simulated networks situated in different dynamical regimes (coloured squares). Eigenspectra of each network accompany each panel, showing bulk distribution (small dots) and outlier (large dot) with respect to the instability limit at unity (dashed line). **B**: Modification of the phase diagram when *N* is fixed (*N* = 1000) and *C* is reduced to increase the degree of sparsity. **C**: Projection of activity along **m** for a network with fixed variance *σ*^2^ and covariance *σ_mn_* (situated at the white square in phase diagrams in **B**) while *N* is fixed (*N* = 1000) and *C* is decreased. The network activity progressively loses structure along **m** since the eigenvalue outlier is reduced. **D**: Same as **C**, but with *C* fixed (*C* = 600) and *N* increased. The outlier is independent of *N*, so structured dynamics can be maintained.

### Input-driven dynamics

For dense rank-one connectivity, when both the eigenvalue outlier and the radius of the bulk are fixed below unity, the fixed point at zero is stable, and the network can display only transient dynamics invoked by an external input *u*(*t*) along a feedforward input pattern **I** (Fig. 4A). If the input pattern is orthogonal to the left connectivity vector **n**, the network activity simply propagates the feedforward input pattern along the one-dimensional axis of **I**. However, two-dimensional trajectories can emerge if the vector **I** is given a non-zero overlap with **n**; in this case, the component of the input-driven activity *ϕ*(**x**) along **n** results in a non-zero *κ_r_*, which allows the trajectory to evolve into the **m** dimension during the course of the input current [8]. This behaviour can be seen by projecting the activity into the **m**-**I** plane (Fig. 4B). The input-driven trajectory is confined to the **m**-**I** plane, revealing the underlying rank-one structure in the connectivity.

To understand the manner in which sparsity interferes with these input-driven trajectories, we first consider what happens when we simply add a random, Gaussian component of variance *g*^2^/*N* to an otherwise dense rank-one matrix, which introduces to the eigenspectrum an eigenvalue disk similar in nature to the bulk distribution that arises under sparsity. With such an addition, the input-driven trajectories are subtly modified as the strength *g* of the random component (Eq. 2) is increased from 0 to 1 (Fig. 4B, C). The projection of the population activity in the **m** -**I** plane remains unaffected (Fig 4B, left), but the random perturbation to the recurrent inputs causes the population activity to gain additional dimensions (Fig. 4B, right). The dominant dimensions of the activity remain aligned with the axes of **I** and **m**, and the degree of activation along the **m** dimension, as quantified by *κ_r_*, is not reduced (Fig. 4C, lower). However, the dimensionality of the network activity increases with *g* (Fig 4C, upper).

The impact of sparsity is markedly different (Fig. 4D-F). As the degree of sparsity is increased, the increased presence of zeros in the connectivity reduces the overlap of the input-driven activity with the **n** dimension, reducing *κ_r_* and leading to a progressive loss of structure along **m** (Fig. 4E, F). Since the feedforward connections are left untouched, the degree to which the activity spans the **I** dimension is not affected. The input-driven trajectories of the sparse network are therefore flattened towards the **I** axis as sparsity is increased (Fig 4E), and the degree of activity along **m** is progressively decreased to zero (Fig 4F, lower). Moreover, despite an initial increase brought about by the full-rank perturbation to the connectivity, the dimensionality of the population activity is ultimately decreased as the gradual pruning of connections weakens the influence of the recurrent inputs (Fig 4F, upper).

We therefore highlight a key difference between the input-driven dynamics of sparsified rank-one networks and those of a dense rank-one network with an added random component. Sparsity interferes with the structure of input-induced activity in a way that a random component does not: it reduces the component of the dynamics along **m** otherwise revealed by an appropriate geometric configuration of inputs, and reduces the overall dimensionality of the activity by stripping away the influence of recurrent inputs. In contrast, increasing the random connectivity in dense networks preserves the dynamics along **m**, and increases the overall dimensionality of the dynamics.

### Autonomous dynamics

When the outlier and the bulk radius in the eigenspectrum are increased above unity, the fixed point at zero loses stability and non-trivial autonomous dynamics emerge. It is here that the equivalence of the sparse regime to the addition of a random component manifests itself. In previous work [8] investigating the autonomous dynamics that emerge in networks comprised of a rank-one component **P** plus a full-rank random part, distinct dynamical regimes were identified on the basis of the dominance of each component in the eigenspectrum. The dynamics were described as *decaying*, if both the outlier of **P** and the eigenvalue disk of **J** lie below unity; *structured stationary*, if the outlier crosses the instability and the disk radius, inducing a non-trivial fixed point along the axis of **m**; or *chaotic*, if the radius of the disk belonging to **J** crosses the instability and the outlier, introducing the higher-dimensional fluctuations classically associated with random networks [24].

Due to the similarities in the eigenspectra, our analyses reveal that the autonomous dynamical regimes of sparsified rank-one networks can be mapped directly onto those described above (Fig. 5). The bulk distribution plays a role analogous to the eigenvalue disk of the random part of the connectivity, while the eigenvalue outlier takes the role of the outlier of **P**. In the rank-one network in the high-sparsity regime (Eq. 12), the location of the bulk and the outlier are independent of *N* and thus remain present in the eigenspectrum even at high sparsities, at magnitudes described respectively by 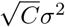 and *Cσ_mn_* (Fig. 3). As in densely connected networks, the instability can either be lead by the outlier, bringing the network into a heterogenous stationary regime aligned with **m** (Fig. 5A, top left), or by the bulk, inducing chaotic dynamics (Fig. 5A, bottom right). Since the magnitude of the bulk radius and the outlier are controlled respectively by the variance *σ*^2^ and covariance *σ_mn_* of the connectivity vectors, the regime in which the network is situated is dictated solely by the relative configuration of these key connectivity parameters. The phase diagram of Fig 5A summarises the dynamical landscape that arises; we note that this diagram is equivalent to that in [8], where the variance *σ*^2^ takes the place of the coupling strength *g* of the random component.

Since the precise location of the outlier and the bulk are a function of *C*, the form of the phase diagram is modulated by *C*. Fixing *N* and modulating *C* shifts the boundaries of the phase diagram (Fig 5B), which can alter the dynamics displayed by a network for given values of the variance *σ*^2^ and covariance *σ_mn_*. For example, in a network fixed at a given parameter location in the phase diagram (white square), progressively decreasing *C* to increase the degree of sparsity causes the network activity to lose structure along **m** (Fig 5C), since the outlier *Cσ_mn_* decreases as a function of *C*.

In contrast, when fixing *C* and increasing *N* to sparsify the connections, the degree of structure along **m** remains unaffected (Fig 5D). This is because the outlier is independent of *N*, and the bulk radius quickly becomes so as the sparsity 1 – *C*/*N* is increased. Thus, the rank-one network constructed as in Eq. 12, appropriately parameterised, can preserve its rank-one outlier and sustain a high degree of sparsity while still displaying the structured, one-dimensional dynamics that are the hallmark of its underlying connectivity.

In summary, sparsified low-rank networks display a wider range of dynamics than their dense counterparts, since the full-rank perturbation to the connectivity introduced by sparsity acts in a manner analogous to the addition of a random component. When the dynamics are purely input-driven, these two cases are not directly equivalent; increasing the degree of sparsity gradually reduces the dimensionality of the dynamics and erodes the structured component, while imposing a random component expands the dimensionality of the dynamics. However, when the dynamics are autonomous, the dynamical regimes accessible to sparsified low-rank networks can be equated directly to those of a rank-one network with an added Gaussian term.

### Computations in sparsified networks

Low-rank recurrent networks benefit from a simple, transparent relationship between the connectivity and resulting dynamics which can be harnessed to implement a rich repertoire of input-output computations [8, 11–13]. The fact that this relationship is preserved even at high levels of sparsity indicates that such computations can be performed even in the highly-sparse regime. By way of example, here we demonstrate how a non-linear input-integration task can be implemented in a sparse low-rank network by designing the connectivity according to intuitive geometric principles originally developed in the context of dense low-rank networks.

We consider a simple evidence-accumulation task in the form of a common behavioural paradigm: a given stimulus is assumed to vary continuously along a particular stimulus feature - such as the coherence of a random-dot kinetogram [28] - and the task is to report whether the magnitude of this stimulus feature is greater than a certain threshold. The actual magnitude is subject to noise fluctuations, so the input must be integrated over time. To model the task, we follow [8] and use the network structure of Fig 6A. We provide the network with a time-varying stimulus *u*(*t*)**I**, where the input pattern **I** represents the stimulus feature of interest and *u*(*t*) is its fluctuating magnitude. A readout unit sums the network activity through a set of readout weights **w** to generate an output *z*(*t*). We require the output to be positive if the stimulus magnitude 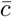 is above threshold, and negative otherwise.

**Fig 6.**
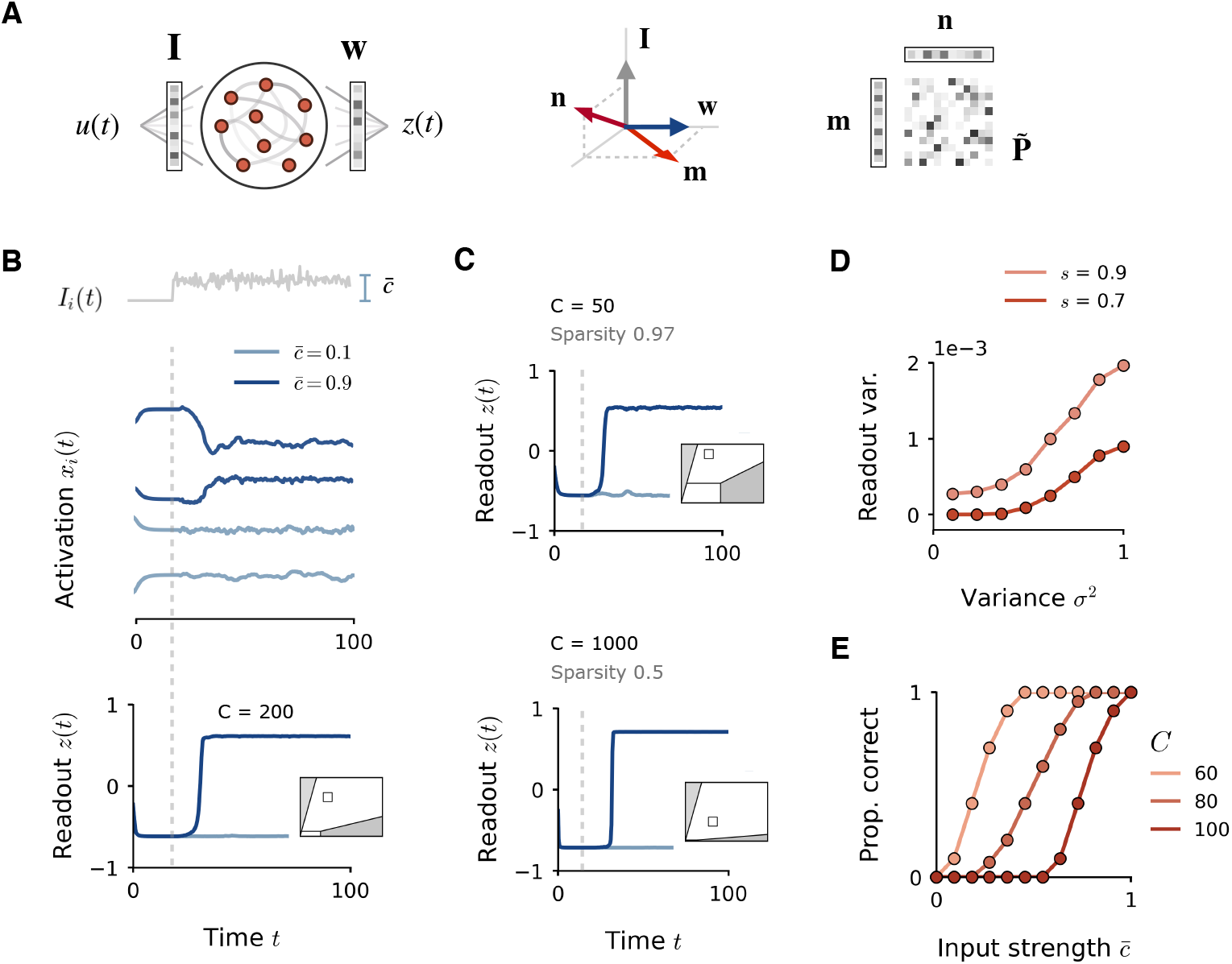
Implementation of input integration task in rank-one networks at high sparsity. **A**: Illustration of recurrent network structure (left), geometrical configuration of input, readout and connectivity vectors (centre), and construction of sparse rank-one connectivity matrix (right). **B**: Implementation of the task in a sparse network (*s* = 0.9, *C* = 200 and *N* = 2000). Top: sample of fluctuating inputs, with magnitude set by coherence parameter 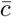. Centre: examples of network activations *x_i_*(*t*) for two inputs of different strengths. Bottom: corresponding readout *z*(*t*) for high and low stimulus strengths. Network is parameterised by variance *σ*^2^ = 0.1 and covariance *σ_mn_* = 0.04, located at the white square on the phase diagram in the inset. **C**: Readout dynamics in networks with differing degrees of sparsity, with *C* modified while *N* is fixed to 2000. Parameters are modified to keep network within the bistable regime (*σ_mn_* = 0.05 and 0.02 respectively), as seen in the phase diagram insets. **D**: Variance in readout response *z*(*t*) to high-coherence inputs 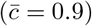 as the variance of the connectivity vectors **m** and **n** is increased. Variance is taken over the final half of the input presentation, for correct responses only. Mean over 50 trials. **E**: Psychometric curve for different sparsity levels *C*, with other parameters held fixed (*σ*^2^ = 0.1, *σ_mn_* = 0.06 and *N* = 2000).

In dense networks, the task can be implemented easily with unit rank connectivity, using a solution which relies solely on an appropriate geometric structure of input and readout weights. The details of the solution are given in [8]; here we simply state the requirements. Firstly, the input pattern **I** must overlap with the left connectivity vector **n**, in order for the input to be picked up by the recurrent dynamics. Secondly, the readout vector **w** must overlap with the right connectivity vector **m**, for the recurrent dynamics to themselves influence the readout. Finally, the connectivity vectors must overlap in a shared dimension orthogonal to the input and the readout, in order to exploit the bistability of the rank-one fixed point and generate the non-linear switch in readout upon integrating the stimulus. To implement the task in a sparse rank-one network, we therefore select connectivity vectors **m** and **n** which obey these requirements (Fig 6A, centre) and construct a rank-one matrix **P** as *P_ij_* = *m_i_n_j_*. We then keep only *C* non-zero inputs per neuron to form the sparse connectivity matrix 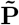 (Fig 6A, right).

Network simulations confirm that the task can be performed accurately at high sparsities (Fig 6B), generating a positive readout only for high stimulus magnitude 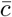. The requirement for success is that the rank-one outlier remains greater than the bulk of the eigenvalue distribution, to keep the network in the structured regime in which the bistability can be maintained. Since the boundaries between regimes shift as *C* is modified (Fig 5B), the connectivity vector overlap *σ_mn_* need only be modulated to ensure that the outlier continues to dominate (Fig 6C, phase diagram insets). If *σ_mn_* is not modulated simultaneously, the psychometric curve for the task simply shifts with *C* (Fig 6E). At high levels of sparsity, the network units receive little recurrent input and are therefore more heavily influenced by noise in the input; however, summing the activations through the readout unit has the effect of averaging away individual fluctuations. As a result, the variation in the readout *z*(*t*) is only mildly increased as sparsity is increased (Fig 6D), and by no means enough to corrupt performance. Likewise, increasing the variance *σ*^2^ of unit-rank connectivity to shift the network towards the chaotic regime has the effect of introducing strong fluctuations into the individual activations, but results in only a small increase in readout variance (Fig 6D), and leaves the task performance unaffected. The implementation of the task is therefore robust to high levels of sparsity and any parameter perturbations which introduce fluctuations to the recurrent dynamics; the key requirement is simply that the bistable structured regime persists, which can be insured by the appropriate relation between input, readout and connectivity vectors.

The basic principle we demonstrate here is that the structured dynamical regime induced by low-rank connectivity can be preserved even at high sparsity, which means that computations which are designed to exploit this structure can be implemented effectively even in networks which are highly sparse. The implication is that the full repertoire of dynamical computations implementable in a low-rank network can be likewise performed at high degrees of sparsity, provided the network is appropriately parameterised.

## Discussion

In this study, we investigated the dynamics of recurrent neural networks in which the connectivity matrices are sparse but possess an underlying low-rank structure. We showed that the resulting full-rank connectivity matrices have eigenspectra which consist of two distinct components, a continuous bulk distribution and isolated outliers. Such eigenspectra are directly analogous to those of a fully-connected unit rank structure plus a full-rank random component [8, 22, 23]. Analytically estimating the magnitude of the outlier and the radius of the continuous bulk in the large *N* limit allowed us to predict the dynamics of the sparsified networks. In particular, the relative magnitude of the two major eigenspectrum components delineates boundaries between decaying, structured and chaotic dynamical regimes. The similarity in the eigenspectra implies that the regimes of autonomous dynamics in sparsified unit-rank networks are analogous to those of dense unit-rank networks with a random connectivity component [8]. Notable differences however appear when the dynamics are purely input-driven. Altogether, we found that computations designed to harness key dynamical properties of low-rank networks are highly robust with respect to sparsity. This identifies sparsified low-rank networks as a biologically-plausible network structure through which to implement computation through low-dimensional population dynamics [7].

The sparse networks examined here were generated by directly removing connections in a fully-connected low-rank structure. The resulting connectivity matrices are directly analogous to those obtained by learning a single pattern through Hebbian plasticity on a sparse subset of connections [9, 29], and therefore are of potential biological relevance. Our analyses can be directly extended to connectivity which consists of a sparsified low-rank structure superposed with a random sparse component with independent entries. As with randomly-connected networks, the results depend on assumptions regarding how the synaptic weights scale with the number of connections [10, 26, 30–33]. Here we considered two cases, and ultimately focused on the situation in which both the number *C* of non-zero connections per neuron and the strength of the connections are fixed as the total number of neurons *N* is increased [26]. Under these assumptions, the radius of continuous bulk of the eigenspectrum remains finite for large *N* [27], as does the value of the outlier. For alternative choices of scaling, our analyses suggest that the expected behaviour of the eigenvalue bulk ultimately depends on the scaling of the variance of the connectivity matrix adjusted by removing the mean low-rank component.

Given its extreme ubiquity in the brain, a question of interest is whether sparsity confers any direct benefit to cortical networks aside from the evident reduction in metabolic and wiring costs. From our analysis, it is not directly clear whether sparsified low-rank networks possess direct computational advantages over their dense counterparts. Insights into the potential computational benefit of sparsity are however rife in the related field of deep learning. Research indicates that the performance of deep networks is remarkably robust to sparsity, and that a large majority of parameters can be pruned without significant loss in accuracy [34, 35]. This makes sparsity a natural regulariser, often employed in combatting over-parameterisation and overfitting [36]. Computational advantages are observed as a consequence, including an improvement in the ability of the trained network to generalise [37, 38] and an increased robustness to adversarial attacks [39, 40], on top of significant savings in memory storage, training time and energy efficiency [35, 41]. Nonetheless, such benefits are most often the result of a highly selective, rule-based pruning process, as opposed to the random weight selection employed in this study. An important avenue of future work will be to explore different forms of structure in the sparsity imposed, and its relation to the training process and the dynamic rules under which the connectivity evolves.

## Methods

### Connectivity vectors

The right and left connectivity vectors **m** and **n** are constructed from three independent normal random vectors **x**, **y** and **z** in the following manner:

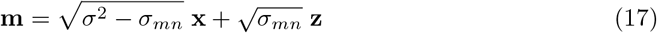

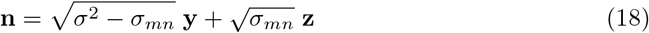

where the components of **x**, **y** and **z** are generated independently from 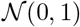. The elements of **m** and **n** are therefore Gaussian-distributed as 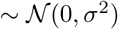 and their degree of overlap onto the **z** direction is controlled by scaling 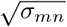 in the interval [0, *σ*].

### Spectral radius of sparsified full-rank matrix

The sparsified full-rank matrix 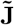 is generated as the elementwise product of the original matrix **J** with an independent random binary matrix **X** whose elements are 0 with probability *s* and 1 with probability 1 – *s*. In other words:

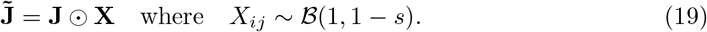

The entries of **J** have a variance of *g*^2^/*N* and a mean of 0, while the entries of **X** have a variance of (1 – *s*). The variance of the entries of 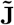 can therefore be derived as:

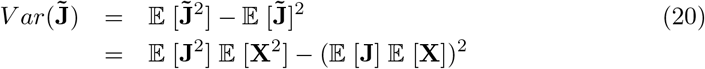

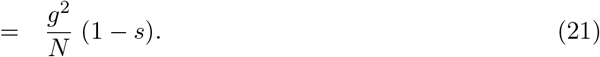

The spectral radius is then given by the circular law:

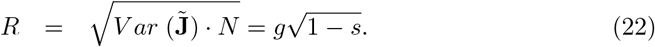

When sparsity is imposed by setting the number of connections per unit *C*, the radius is given by substituting *s* = 1 – *C*/*N* as:

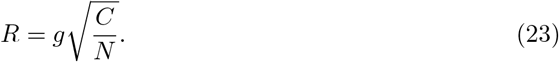

### Eigenvalue spectrum of sparsified rank-one matrix

The elements of the sparsified rank-one matrix 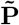 are generated in an equivalent manner as 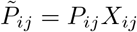. Its eigenspectrum is comprised of a continuous bulk and an isolated outlier. We here derive the location of the outlier λ_1_ and the radius *R* of the bulk distribution individually.

#### Outlier

We proceed by showing that, for *N* → ∞, the right connectivity vector **m** is an eigenvector **v** of 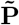, and derive the corresponding eigenvalue λ. By writing the individual matrix elements of 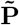 as:

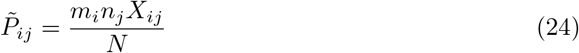

the *i^th^* element of 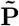 applied to **m** is:

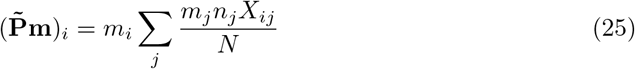

As *N* → ∞, the sum over *j* in the right-hand side converges to an expectation due to the central limit theorem, and we may write:

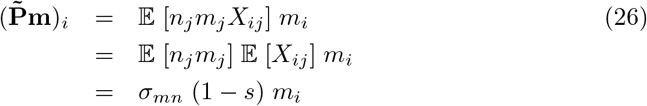

It therefore holds that:

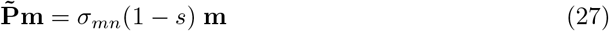

so that **m** is a right eigenvector of 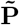 with eigenvalue

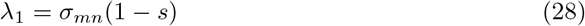

#### Bulk

To determine the radius of the bulk distribution, we derive the variance of the elements of the matrix with the outlier eigenvalue removed, 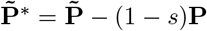. We first rewrite **X** as 1 – **B**, where **B** is a Bernoulli matrix with entries 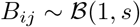, in order to rewrite the individual matrix entries 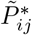, as follows:

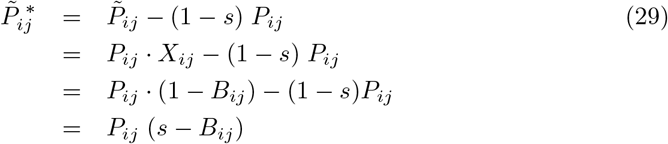

We can then derive the variance of the entries 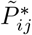, as

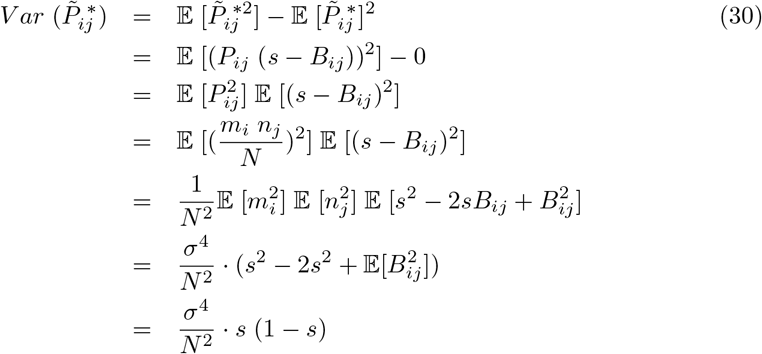

given that 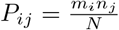, and *m_i_* and *n_i_* each have variance *σ*^2^. As before, we may substitute *s* = 1 – *C*/*N* when sparsity is imposed by setting the number of connections per unit, to give:

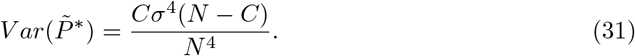

The bulk radius is then obtained via the circular law 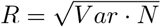 as in the Gaussian case.

### Latent dynamics in low-rank networks

Here we summarize the description of low-dimensional dynamics in low-rank networks [12, 13].

The low-rank network of Eq. 3, with connectivity matrix 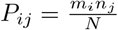, is governed by the dynamics introduced in the main text (Eq. 1):

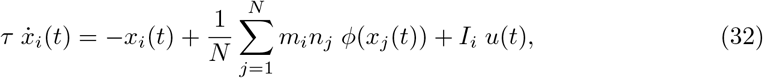

At the level of the population, the collective trajectory **x** (*t*) is embedded in a low-dimensional linear subspace [12, 13]. The total dimensionality of this subspace is the sum of the rank of the connectivity matrix **P** plus the dimensionality of external inputs. For a rank-one network with one external input vector, the dynamics are constrained to the two-dimensional plane spanned by the left connectivity vector **m** and the input vector **I**. The dynamics can then be represented in a new basis by projecting the activity trajectory **x** (*t*) onto these two axes. The individual unit activations thus read as:

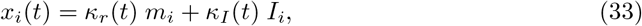

where *κ_r_*(*t*) and *κ_I_*(*t*) are projections of the activity **x**(*t*) onto the **m** and **I** axes respectively:

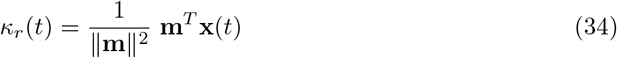

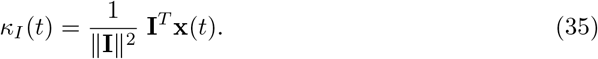

The projection onto the **m** axis, *κ_r_*(*t*), is then governed by its own dynamical equation:

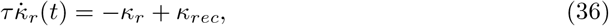

where:

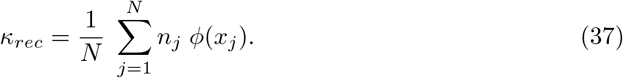

At equilibrium, we therefore have *κ_r_* = *κ_rec_*.

## Acknowledgments

This work was supported by the CRCNS project PIND funded through the National Institute of Health (NIMH: 1R01MH122025-01) and French Agence Nationale de la Recherche (ANR-19-NEUC-0001-01), and the program “Ecoles Universitaires de Recherche” launched by the French Government and implemented by the ANR, with the reference ANR-17-EURE-0017.

